# SARS-CoV-2 Neutralization in Convalescent Plasma and Commercial Lots of Plasma-derived Immunoglobulin

**DOI:** 10.1101/2021.08.13.456066

**Authors:** Andreas Volk, Caroline Covini-Souris, Denis Kuehnel, Christian De Mey, Jürgen Römisch, Torben Schmidt

## Abstract

**Introduction:** Patients suffering from primary or secondary immunodeficiency (PID or SID) face times of increased insecurity and discomfort in the light of the raging COVID-19 pandemic, not knowing if and to what extent their comorbidities impact the course of a potential SARS-CoV-2 infection. Furthermore, recently available vaccination options might not be amenable or effective for all patients of this heterogeneous population. Therefore, these patients often rely on passive immunization with plasma-derived, intravenous or subcutaneous immunoglobulin (IVIG/SCIG).

Whether the ongoing COVID-19 pandemic and/or the progress in vaccination programs lead to increased and potentially protective titers in plasma-derived immunoglobulins (Ig) indicated, e.g., for humoral immunodeficiency remains a pressing question for this patient population.

**Purpose:** We investigated SARS-CoV-2 reactivity of US plasma-derived IVIG/SCIG products from the end of 2020 until 06/2021 as well as in convalescent plasma (CP) from 05/2020 to 08/2020 to determine whether potentially neutralizing antibody titers may be present.

**Methods:** Final containers of IVIG/SCIG and CP donations were analyzed by commercial ELISA for anti-SARS-CoV-2 S1-receptor binding domain (RBD) IgG as well as microneutralization assay using a patient-derived SARS-CoV-2 (D614G) isolate. Neutralization capacities of 313 plasma single donations and 119 plasma-derived IVIG/SCIG lots were determined. Results obtained from both analytical methods were normalized against the WHO International Standard. Finally, based on dense pharmacokinetic (PK) profiles of an IVIG preparation from previously published investigations, possible steady-state plasma levels of SARS-CoV-2 neutralization capacities were approximated based on currently measured anti-SARS-CoV-2 potencies in IVIG/SCIG preparations.

**Results:** CP donations presented with a high variability with regards to anti-SARS-CoV-2 reactivity in ELISA as well as in neutralization testing. While approximately 50% of convalescent donations were none/low neutralizing, approximately 10% were at or above 1000 IU/mL.

IVIG/SCIG lots derived from pre-pandemic plasma donations did not show neutralizing capacities of SARS-CoV-2. Lots produced between 12/2020 and 06/2021, entailing plasma donations after the emergence of SARS-CoV-2 showed a rapid and constant increase in anti-SARS-CoV-2 reactivity and neutralization capacity over time. While lot-to-lot variability was substantial, neutralization capacity increased from a mean of 20 IU/mL in 12/2020 to 505 IU/mL in 06/2021 with a maximum of 864 IU/mL for the most recent lots.

Pharmacokinetic extrapolations, based on non-compartmental superposition principles using steady-state reference profiles from previously published PK investigations on IVIG in PID, yielded potential steady-state trough plasma levels of 16 IU/mL of neutralizing SARS-CoV-2 IgG based on the average final container concentration from 05/2021 of 216 IU/mL. Maximum extrapolated trough levels could reach 64 IU/mL based on the latest maximal final container potency tested in 06/2021.

**Conclusions:** SARS-CoV-2 reactivity and neutralization capacity in IVIG/SCIG produced from US plasma rapidly and in part exponentially increased in the first half of 2021. The observed increase of final container potencies is likely trailing the serological status of the US donor population in terms of COVID-19 convalescence and vaccination by at least 5 months due to production lead times and should in principle continue at least until fall 2021. In summary, the data support rapidly increasing levels of anti-SARS-CoV-2 antibodies in IVIG/SCIG products implicating that a certain level of protection could be possible against COVID-19 for regularly substituted PID/SID patients. Nevertheless, more research is still needed to confirm which plasma levels are needed to provide protection against SARS-CoV-2 infection in immune-compromised patients.

**Plain Language Summary:** People with deficiencies in their immune system often have an insufficient antibody response to antigens, e.g., bacteria, viruses, or vaccines. These patients therefore often receive antibodies from healthy people to replace the missing antibodies and build a first line of defense against infections. These antibodies (also called immunoglobulins (Ig)) are prepared from plasma of healthy donors, the liquid fraction of the blood without cells. This plasma is then split up in pharmaceutical production into its protein components. One of these is immunoglobulin G (IgG), which is the protein family that neutralizes/inactivates infectious agents as well as marks these infectious agents so they can be recognized by other parts of the immune system. With the ongoing COVID-19 pandemic and the severe to fatal outcomes for certain patient groups, especially people with impaired immunity, these patients and their physicians are interested in whether their antibody replacement therapy also confers protection against SARS-CoV-2 infection. We analyzed the capability of plasma-derived Ig lots to (i) recognize SARS-CoV-2 protein by ELISA method as well as (ii) neutralize SARS-CoV-2 by neutralization studies using the actual virus under biosafety level 3 (BSL-3) conditions. Here we show increasing anti-SARS-CoV-2 activity over time of manufactured Ig lots produced between 12/2020 and 06/2021. The most recent lots had a neutralizing activity of up to 864 IU/mL. Considering that the USA represents Octapharma’s main plasma source, the progress in vaccination levels together with the evolution of the COVID-19 pandemic in this country suggests that the IVIG/SCIG neutralization capacities against SARS-CoV-2 might still increase and could potentially meet a level where antibody plasma concentrations in the patient confer immune protection.

**Key Points:**

- Patients with humoral immunodeficiency rely on plasma-derived immunoglobulin for passive immunization against numerous pathogens.
- SARS-CoV-2 neutralization capacities of plasma-derived immunoglobulins have increased over time with the ongoing COVID-19 pandemic and vaccination campaigns.
- Plasma-derived immunoglobulin in prophylactic use for immunodeficient patients could potentially protect against SARS-CoV-2 infection in the future.

## 1. Introduction

The principle of passive immunization, i.e., the infusion of antibodies from a healthy immunized or convalescent donor to patients not mounting their own active immune response, is a long-standing treatment option for infectious diseases and was first applied by Emil von Behring, who received the first ever Nobel prize in 1901 for serum therapy of diphtheria [1]. Passive immunization refers to antibody transfer by means of recombinant monoclonal antibody, plasma-derived intravenous or subcutaneous immunoglobulin (IVIG/SCIG), or convalescent plasma (CP). In fact, CP therapy has faced a renaissance as a possible treatment option in the currently ongoing COVID-19 pandemic to bridge the period until therapeutics, and especially vaccines as protective measures, became available [2–4]. FDA granted emergency use authorization for CP therapy in 08/2020 and revised this guidance in 03/2021 to use only higher titer CP early in the course of disease [5]. However, for the therapeutic setting a meta-analysis by Peng et al. identified a total of 243 published studies including 64 clinical trials on CP administration for prevention and treatment of COVID-19 by 04/2021 [6]. In summary, the efficacy of CP therapy against COVID-19 has not been unequivocally shown and remains a matter of debate and further clinical investigation [7–12]. The criticality of early treatment with CP as well as the required dose and specific antibody titers of the donations have been discussed as part of relevant drivers for treatment success [11]. Likewise, the prophylactic use of CP has not found broad application [6]. However, efficacious prophylaxis against disease using CP has been shown in hamster and macaque models [13–15]. Furthermore, clinical efficacy has been shown with recombinant monoclonal antibody prophylaxis supporting the notion that immunoglobulin G (IgG) molecules can prevent infection/COVID-19 [16, 17].

Active immunization of patients with primary immunodeficiencies (PID) as well as secondary immunodeficiencies (SID), which in part manifest as impeded or a complete lack of humoral response, has recently been reported with encouraging findings [18, 19]. However, not all patients within this heterogeneous group are amenable to vaccination or vaccination might not lead to immune protection [20, 21]. Furthermore, COVID-19 disease in this indication group has been reported to have both less [22] and more severe outcomes depending on comorbidities and individual patient factors [23].

Regardless of the amenability of active vaccination and the potential disease outcome, passive immunization by repeated injection of IVIG/SCIG for at least a part of this patient group is the standard of care for humoral deficiency and has been ongoing prior to, during, and after the COVID-19 pandemic [24–26]. IVIG/SCIG are purified and concentrated immunoglobulin (Ig) preparations derived from pooled plasma donations [27, 28]. Therefore, the reactivity and neutralizing potency of these IgG pharmaceutical products are directly dependent on the epidemiologic experience of the donor population [19–21].

It is thus of high interest - in the light of increasing seroconversion in the donor population - how currently manufactured IVIG/SCIG can serve these immunodeficient patient groups in terms of protection against a SARS-CoV-2 infection or impede disease severity of COVID-19. First reports of anti-SARS-CoV-2 reactivity and neutralization capacities in commercially produced immunoglobulins have already been published [29–33].

To obtain an impression on how CP donations vary in terms of potency, we first investigated a selection of COVID-19 CP donations from 05/2020 to 08/2020 for SARS-CoV-2 reactivity by use of commercially available anti-SARS-CoV-2 IgG ELISA and an inhouse established SARS-CoV-2 microneutralization assay under biosafety level 3 (BSL-3) conditions to assess the actual neutralization capacity.

Furthermore, we analyzed commercial lots of immunoglobulin manufactured from 11/2020 to 06/2021 by ELISA. A fraction of these lots was subsequently subjected to neutralization testing against actual SARS-CoV-2 virus.

Lastly, we performed calculations based on previous pharmacokinetic (PK) profiles of IVIG [34] to obtain a first and preliminary insight into possibly achievable steady-state plasma levels of anti-SARS-CoV-2 IgG in patients regularly dosed with IVIG.

## 2. Material and Methods

### 2.1. Single Donation of Convalescent Plasma

Analyzed CP donations were collected from 05/2020 to 08/2020. Octapharma invited donors to present proof of resolved SARS-CoV-2 infection (either by positive diagnostic test or positive serological test) to be eligible to donate CP after a deferral period of 14 days after either diagnostic test or symptom cessation, as applicable.

### 2.2. IVIG/SCIG

Octagam^®^ and Panzyga^®^ are polyclonal IVIG and Cutaquig^®^ is a polyclonal SCIG product derived from thousands of plasma donations. We tested lots derived from US donations prepared from 11/2020 to 06/2021. Octagam^®^ is available as 50 mg/mL and 100 mg/mL (5% and 10%) preparations, Cutaquig^®^ is available as 165 mg/mL (16.5%) while Panzyga^®^ is available as 100 mg/mL (10%) IgG preparations. All reported results were normalized to a potency of 100 mg IgG/mL to remove bias from different formulations.

### 2.3. Anti-SARS-CoV-2 IgG ELISA Kit

The following ELISA kits were used for detection of anti-SARS-CoV-2 IgG against receptor binding domain (RBD) of spike protein S1: EI-2606-9601 G ELISA kit (EUROIMMUN Medizinische Labordiagnostika AG, Lübeck, Germany) for qualitative detection; QuantiVac EI-2606-9601-G ELISA (EUROIMMUN Medizinische Labordiagnostika AG, Lübeck, Germany) for quantitative determination of IgG titers. Tests were performed as instructed by the provider.

CP was stored frozen and thawed promptly before analysis. Samples were diluted with buffer solutions provided in each ELISA kit. After qualitative detection, results were expressed as antibody ratio.

Concentrated IVIG/SCIG lots were diluted using buffer solutions included in each ELISA kit. 5%, 10%, and 16.5 % IgG preparations were tested in 3 dilution series ranging from 1:1 to 1:10. All samples were tested in 8 replicates. Results were initially obtained in relative units (RU/mL) and were normalized against the First WHO International Standard (NIBSC code: 20/136) in binding antibody units/mL (BAU/mL) as described by the test kit provider. All quantitative results were normalized to 100 mg/mL IgG for comparison.

### 2.4. Cell Culture

Vero cells (CCL-81, American Type Culture Collection) were cultured in RPMI-1640 medium (Sigma^®^), supplemented with 10% fetal bovine serum (Sigma^®^) and 1% penicillin/streptomycin (Sigma^®^) at 37°C, 5% CO_2_ in saturated humidity.

FRhK-4 cells (CRL-1688, American Type Culture Collection) were cultured in EMEM medium (Sigma^®^) supplemented with 14% RPMI-1640 (Sigma^®^), 5% fetal bovine serum and 1% penicillin/streptomycin (Sigma^®^) at 37°C, 5% CO_2_ in saturated humidity.

### 2.5. Virus

SARS-CoV-2 D614G (Human 2019-nCoV ex China_BavPat1/2020_Germany ex China, GISAID ID: EPI_ISL_406862) was kindly provided by Prof. Dr. Christian Drosten, Institute of Virology, Charité, Universitätsmedizin Berlin, Germany [35].

All handling of virus was performed under BSL-3 conditions according to German law and regulations.

The virus was propagated on FRhK-4 cells established three days prior to inoculation with 20 mL of 21.5 × 10^3^ cells/mL in FRhK-4 culture medium (see above) in a T75 cell culture flask (Falcon^®^). Three days later, the culture medium was removed completely. Cells were inoculated with SARS-CoV-2 at a multiplicity of infection of 0.01 in a total of 5 mL virus suspension. Cell inoculation was incubated for 1 h at 37°C, 5% CO_2_ in saturated humidity. Inoculum was removed and the cell layer was washed once with 1x phosphate-buffered saline (Sigma^®^). After addition of 30 mL FRhK-4 culture medium, cells were maintained for 4 days. Subsequently, the medium was exchanged to FRhK-4 medium without fetal bovine serum. On day 6 after inoculation, virus-containing supernatant was removed and centrifuged to remove cells and cellular debris (4°C, 15 min at 1500x g). The supernatant was kept refrigerated during cell lysis steps. The cell monolayer in the T75 culture flask as well as the centrifugal pellet were frozen three times in −80°C ultra-deep freezer for cell lysis. The flask was then rinsed thoroughly with 5 mL of the refrigerated virus-containing supernatant and subsequently used to resuspend the freeze-thawed centrifugal pellet. After repeated centrifugation of the cell lysate suspension, the supernatant was added to the remainder refrigerated virus suspension followed by 0.45 μm filtration. After aliquoting, virus was stored in a −80°C ultra-deep freezer until use.

### 2.6. Standards and Reagents

#### 2.6.1. Neutralization Standards

WHO International Standard / First WHO International Standard for anti-SARS-CoV-2 immunoglobulin (human) was obtained from the National Institute for Biological Standards and Control (NIBSC), Potters Bar, Hertfordshire, EN6 3QG, UK, NIBSC code: 20/136 [36]. The WHO International Standard has a rated potency of 1000 IU/mL or 1000 BAU/mL depending on assay type, if reconstituted according to instructions.

The internal neutralization standard “V” was generated in-house at Octapharma Virus and Prion Validation by pooling a selection of CP samples. After pooling, samples were aliquoted and kept frozen at −80°C until use. Standard was diluted 1:40 initial dilution in 20% citrate dextrose (ACD, Sigma^®^) in Vero culture medium to avoid clotting.

The internal neutralization standard “O” was a high-titer, single source plasma donation obtained from Octapharma Plasma Inc., Charlotte, NC, USA. After thawing, the donation was aliquoted and kept frozen at −80°C until use. Standard was diluted 1:40 initial dilution in 20% ACD in Vero culture medium to avoid clotting.

### 2.7. Neutralization Testing

Neutralization testing (NT) was done in two microneutralization assay formats differing in sample dilution schemes and replicates. Both assays were not prospectively validated. In both, Vero cultures were prepared one day prior to assay conduct and 120 μl was seeded in each of the 96 wells with 0.5×10^4^ cells/mL (= 600 cells per well).

In the screening assay format, which was used in the CP donation testing, each plasma sample was thawed at 37°C and initially diluted 1:40 with 20% ACD in cell culture medium to avoid clotting. Subsequently, samples were serially diluted 1:2-fold in six consecutive steps. Each of the four parallel dilutions was assayed once for neutralization. The assay principle was adapted from Gauger & Vincent [37].

In the potency assay format deployed for final container testing, each sample was assayed in 3 independent dilution series. After an initial 1:10 dilution step, 10 consecutive 1:2 dilution steps were performed. Each dilution was assayed in octuplicates, respectively. This assay format has been previously published [30, 38]. It shows more reliable results per sample, although throughput is limited and is also restricted owing to BSL-3 procedures. Experimental performance of both screening and potency assay formats are summarized in Supp. Fig. 1.

In both assay versions, each sample dilution was subsequently challenged with an equal volume of SARS-CoV-2 suspension, which had a titer of 3.0 log_10_ TCID_50_/mL. The virus titer was confirmed in each experiment run by so-called back titrations (refer to section “virus titration”). A mean titer of n=5 titrations was determined to accurately account for the actual viral load in the immune neutralization, which was used as a normalizing factor to account for potential inter-run variances.

In both formats, the neutralization reaction was incubated for 150 ± 15 min at 37°C in a CO_2_ incubator at saturated humidity. After the incubation period, 100 μl of each replicate dilution was transferred onto a 96-well culture of Vero cells (see above). Inoculated cultures were maintained for 7 days at 37°C, 5% CO_2_ in saturated humidity. On day 7, cultures were microscopically evaluated for the presence or absence of cytopathic effect by two independent operators (four eyes-principle). Neutralization potency by means of dilution to achieve a 50% neutralization of virus was calculated using Spearman-Kärber-based statistics [38].

To convert the identified NC50 dilution values into international units, the normalized (mean) neutralization values of the sample were divided by the (mean) neutralization of the internal standard run along the same experiment and then multiplied by the calibrated potency of the internal standard in international units.

### 2.8. Virus Titration

On each day of neutralization, the virus stock used as a challenge inoculum for neutralization studies was assessed via repeated titration by means of Spearman-Kärber statistics [39]. For that, a serial 1:3 dilution of the virus stock was prepared in 12 consecutive steps in Vero cell culture medium. Subsequently, 8 × 100 μl of each dilution were added to a 96-well plate containing Vero cultures established one day earlier as described above. Inoculated cultures were maintained for 7 days at 37°C, 5% CO_2_ in saturated humidity. On day 7, cultures were microscopically evaluated for the presence or absence of cytopathic effect by two independent operators.

### 2.9. Pharmacokinetic Analysis

Steady-state levels of SARS-CoV-2 neutralizing IgG on 4-weekly dosing were approximated by superposition principles using baseline-adjusted, dose-normalized reference profiles from a previously published clinical trial (EudraCT 2009-011434-10) that investigated the steady-state pharmacokinetics of IgG on 4-weekly repeated dosing of IVIG 10% in patients with PID [34]. The individual courses of total IgG were baseline-adjusted, assuming that the baseline levels were residuals from previous dosing whilst endogenous levels were negligible. To be able to pool the profiles across subjects, the baseline-adjusted profiles were dose-normalized. The time courses of the median, 25^th^ (P25), and 75^th^ (P75) percentile levels of the baseline-adjusted dose-normalized data were used as reference profiles. These reference profiles were expanded by linear/log-linear interpolation between sampling points and extrapolation beyond the last quantifiable value by means of the apparent terminal disposition rate constant. Steady-state levels were approximated by building time-staggered superposition cascades of the reference profiles for ten 4-weekly doses, assuming the PK to be dose-proportional and time-invariant.

### 2.10. Statistical Methods

All statistical methods are identified in the respective figure legends. For statistical analyses, the software GraphPad Prism version 8.4.3 (686) for Windows, GraphPad Software, San Diego, California USA, www.graphpad.com, was used.

PK analyses were carried out by means of PCModfit v7.0 (2021); Copyright of Graham D. Allen, available at https://pcmodfit.co.uk/index.html, accessed on 23.07.2021.

The minimal serorelevant exposure in % of the US population (as depicted in Fig. 2b) was calculated by data obtained from CDC Covid Tracker [40] as follows: The sum of “Cumulative Count of People Receiving at least One Dose Reported to CDC by Date Administered, United States” (derived from https://covid.cdc.gov/covid-data-tracker/#vaccination-trends_vacctrends-onedose-cum” accessed on 12.07.2021) and the “Trends in Total [cumulative] Cases in The United States Reported to CDC” (derived from https://covid.cdc.gov/covid-data-tracker/#trends_totalcasescumulative accessed on 12.07.2021). This sum was the divided by the total US population (derived from https://www.census.gov/popclock/; accessed on 02.06.2021) [41].

## 3. Results

We obtained CP donation units from Octapharma Plasma’s donation centers in the US between 05/2020 and 08/2020 to understand early in the pandemic how CP donations present themselves in terms of titer and inter-donation variation. We therefore first tested 133 single donations by means of qualitative anti-SARS-CoV-2-S1 RBD IgG ELISA. We found a positive signal of anti-SARS-CoV-2 IgG in 79% of CP donations, while 15% and 6% of the donations exhibited borderline and negative results, respectively. The antibody ratio ranged from <0.1 (limit of detection, LOD) up to 16 while the mean ± standard deviation (SD) ratio was 4.5 ± 3.6 (Fig. 1a).

**Fig. 1.**
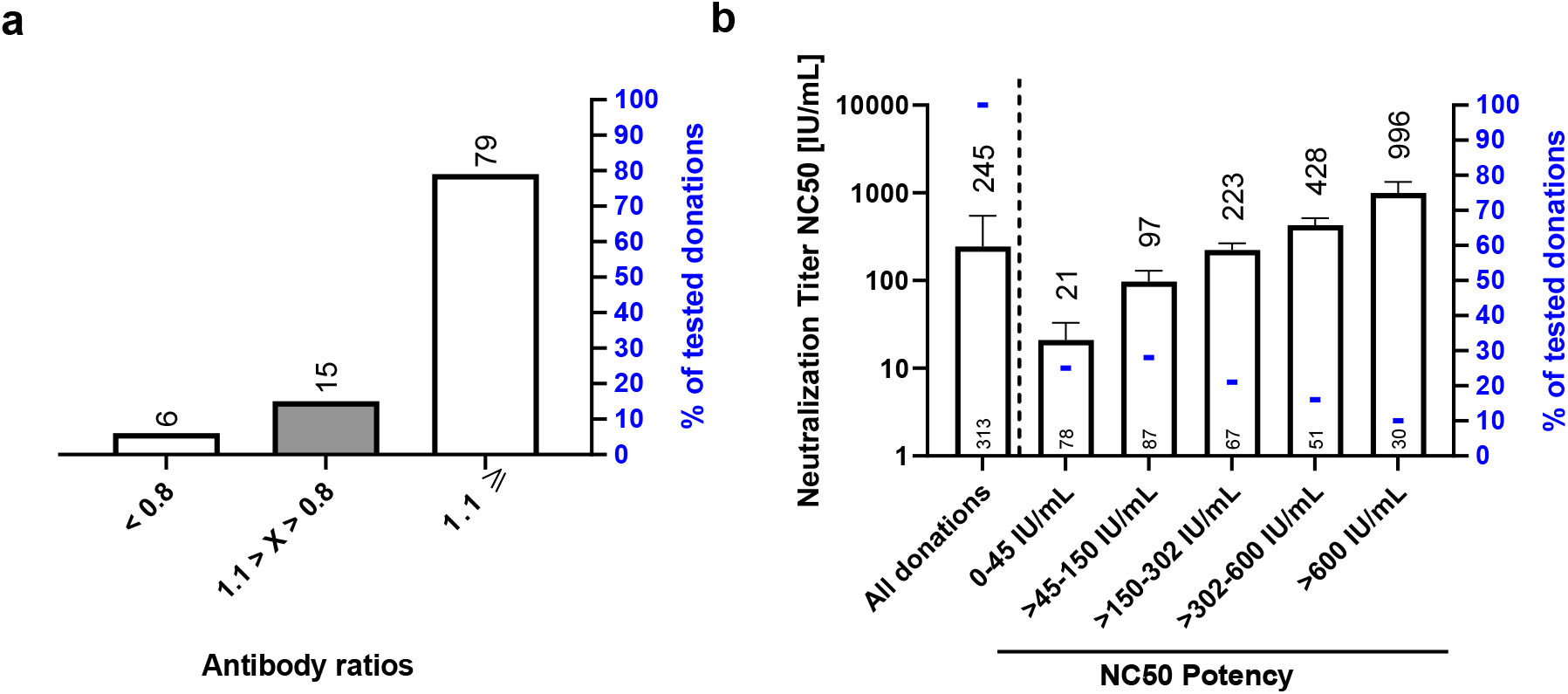
Characterization of CP single donations for anti-SARS-CoV-2 reactivity by means of **a** IgG ELISA and **b** microneutralization assay. **a** ELISA ratios of anti-SARS-CoV-2 IgG are plotted. Cut-off is set by ELISA kit manufacturer as 1.1. Interpretation was negative if ratio is < 0.8; positive if ≥ 1.1; borderline if 1.1 > x > 0.8. **b** Samples are clustered into neutralization potency (NC50) bins (right) and also shown as total sample set (left). For each ranked sub-population, the mean normalized neutralization titers ± SD are depicted on the left y-axis and values are shown above bars. The number of donations binned to each NC50 potency subset is shown at the bottom of each bar. The fraction of each subset to the total sample size is indicated by blue horizontal lines and depicted on the right y-axis. IU: international units

We next set out to investigate what these ELISA data mean in terms of actual virus neutralization. Therefore, 313 CP donations including the above sample set were analyzed by actual virus neutralization in a so-called NT screening assay format (Fig. 1b). We found a high variability of neutralization activity among CP donations. 25% had a neutralization capacity close to or below the LOD (0-45 IU/mL). 28% had only a weak neutralization titer of less than 150 IU/mL. 21% of donations had a mean titer of 223 IU/mL, representing a moderate/average neutralization. 16% had a neutralization titer between 300 IU/mL – 600 IU/ml, while only 10% of all donations had a neutralizing activity of at least 600 IU/mL. The maximal donation neutralization titer found was 2017 IU/mL. The mean ± SD neutralization titer of all convalescent donations was 245 ± 302 IU/mL. Despite this high variability, we could verify a mean convalescent level of around 200 IU/mL in other CP donation cohorts (data not shown).

Given the reports on frequent asymptomatic to mild COVID-19 infections and the general interest on how epidemiologic developments (infection as well as vaccination programs) in the US donor population reflects in IVIG/SCIG, we investigated commercial lots derived from US plasma sources for SARS-CoV-2 reactivity from 11/2020 until 06/2021 by means of quantitative anti-SARS-CoV-2 S1 RBD IgG ELISA. (Fig. 2a). Pre-pandemic lots all tested negative in the ELISA, excluding cross-reactivity of the assay against ‘common-cold’ coronaviruses. Systematic investigations started in 11/2020 and the first reactive lots were detected in 12/2020 with ELISA IgG titers ranging from 40 to 170 BAU/mL with a mean value of 100 BAU/mL. A progressive reactivity increase was observed, which took up pace in the very last months of observation. A mean ± SD anti-SARS-CoV-2 IgG titer of 1048 ± 363 BAU/mL was found in IVIG/SCIG lots produced in 06/2021.

**Fig. 2.**
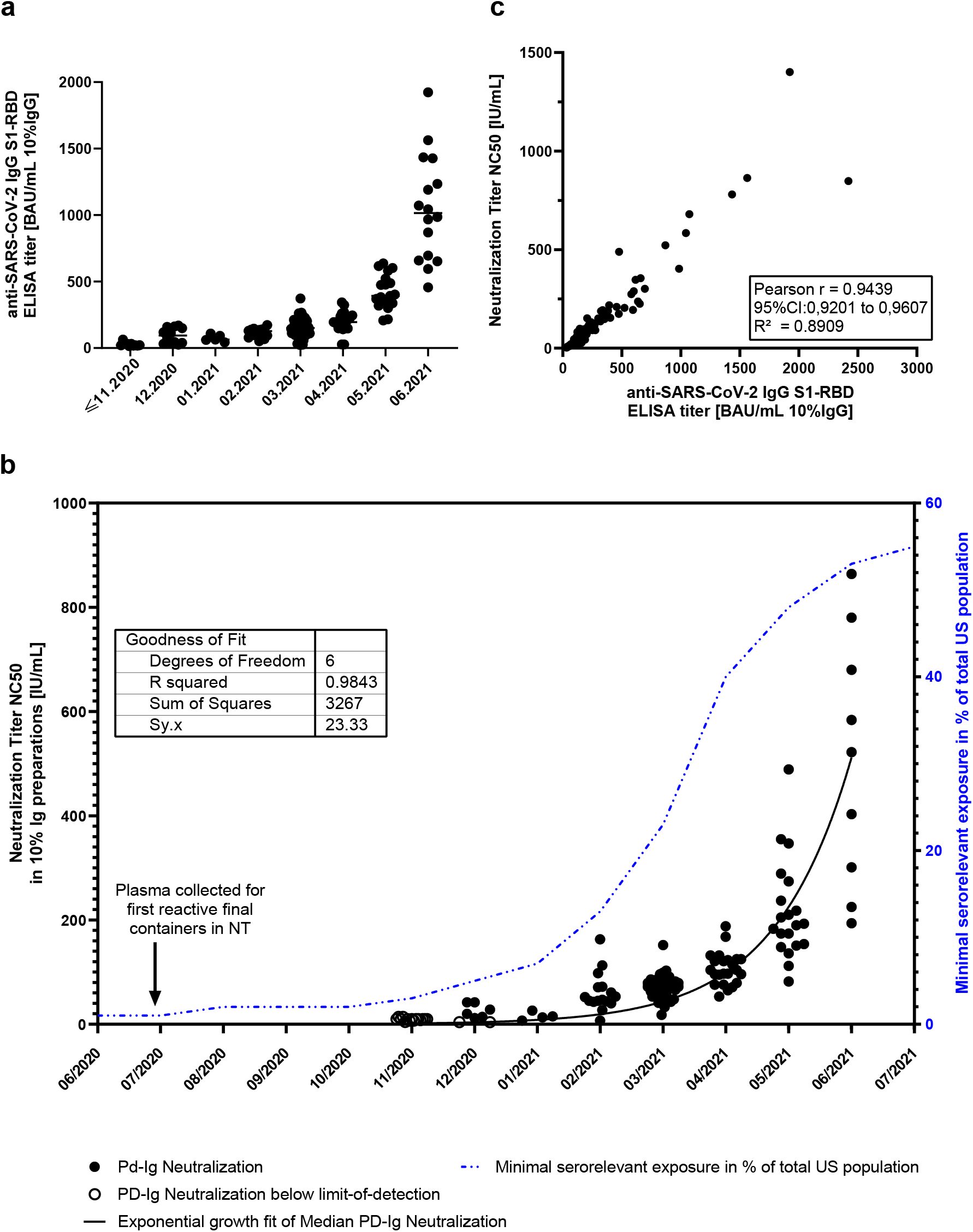
SARS-CoV-2 reactivity and neutralization titer of IVIG/SCIG final containers from 11/2020 – 06/2021. **a** Scatter plot for reactivity of 10% IVIG/SCIG final containers normalized to 10% Ig in commercial IgG ELISA. Each dot represents the mean result of 8 determinations per sample counted as one measurement. The week of production is plotted against ELISA signal intensity, which was normalized against the WHO International Standard and to 10% IgG (100 mg/mL). **b** Neutralization titer of 10% IVIG/SCIG as determined by microneutralization assay is depicted on the left y-axis in IU/mL. Each dot represents the mean of triplicate analysis, counting as one measurement. Neutralization over time was fitted (solid line) using non-linear fit “exponential growth equation” on the median values under GraphPad Prism version 8.4.3 (686). The cumulative minimal serologically relevant exposure of the total US population is depicted as dash-dotted line (right y-axis) in blue. The latter were obtained from CDC Covid Data Tracker [40]. **c** Correlation analysis of ELISA (x axis) and microneutralization results (y-axis). Correlation coefficient was calculated using two-tailed, Gaussian distributed values at 95% CI (Pearson) under GraphPad Prism version 8.4.3 (686). BAU/mL: Binding antibody units

**Fig. 3.**
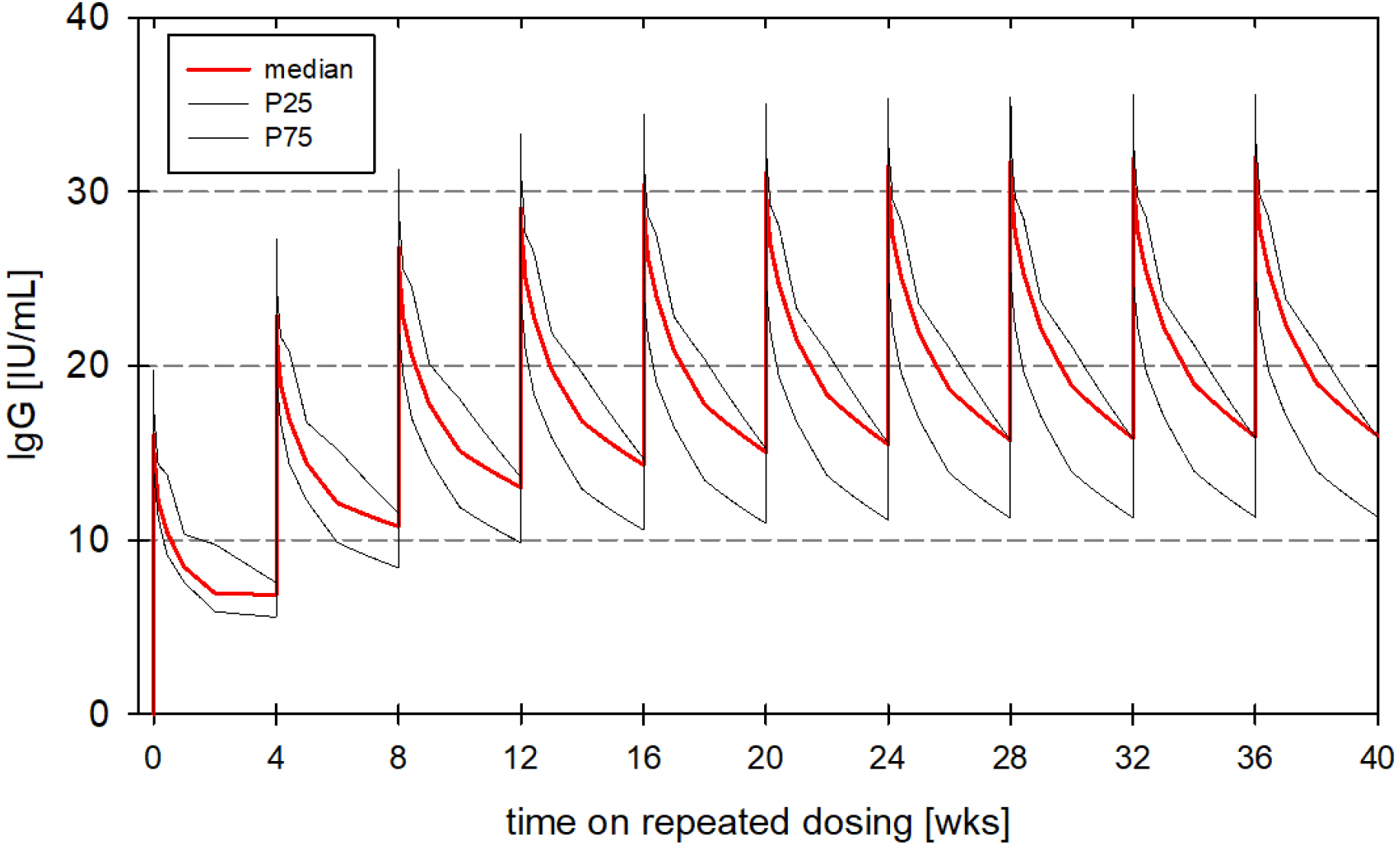
Approximated time courses of the neutralizing anti-SARS-CoV-2 IgG-levels (IU/mL) throughout ten 4-weekly repeated doses of intravenous IgG (potency 216 IU/100 mg) dosed with the median dose of the reference dataset (0.406 g/kg-BW). Data was processed using PCModfit v7.0 (2021). PK data was obtained from a previously published study [34]

To substantiate this information, we further tested ELISA-positive lots in NT in the potency assay format, (Fig. 2b).

As was seen with the ELISA data, the first neutralizing IVIG/SCIG lots were identified in 12/2020 (mean ± SD neutralization titer of 21 ± 16 IU/mL). An exponential, yet slow increase in neutralization activities was subsequently observed (i.e., mean values ± SD in 02/2021: 61 ± 37 IU/mL; 04/2021: 107 ± 32 IU/mL; 05/2021: 216 ± 96 IU/mL; and 06/2021: 506 ± 242 IU/mL). The maximum NC50 measured was 864 IU/mL. We fitted the median SARS-CoV-2 neutralizing titers by exponential growth equation fit (least squares regression without weighing, no outlier handling and constraints) and found an R^2^ of 0.98.

For comparison and as a possible reference to the US plasma donor population, data adapted from CDC epidemiologic surveillance on the relative minimal SARS-CoV-2 exposure of the total US population [%] is referenced in Fig. 2b [40].

We also performed correlation of the anti-SARS-CoV-2 IgG ELISA data and the NT results and found a correlation coefficient of r = 0.94 (Pearson, two-tailed, 95% CI) with p<0.0001 indicating a quantitative character of this IgG ELISA. The linear regression R^2^ was 0.89 (Fig. 2c).

Finally, we investigated how the currently measured final container potencies relate to steady-state plasma levels of anti-SARS-CoV-2 IgG in a patient receiving IVIG 10%. We deployed data from a previously published trial (EudraCT 2009-011434-10) with PK characterization of the levels of total IgG on repeated 4-weekly IVIG dosing (median dose: 27.4 g, inter-quartile range (IQR): 20.4 g to 29.1 g) in 30 patients (11 females, 19 males; 8 children, 4 adolescents, and 18 adults) with PID (median body weight (BW): 67.7 kg, IQR: 52.4 kg to 75.3 kg) [34]. As detailed in the methods section, individual time courses of the untransformed IgG levels (Supp. Fig. 2a) were baseline adjusted (Supp. Fig. 2b) and subsequently normalized to the IVIG dose administered (Supp. Fig. 2c). Ultimately, the normalized, baseline adjusted one-dose PK profiles were expanded by staggered superposition to achieve an assumption of a PK time-course of a 4-weekly IVIG treatment regimen (Supp. Fig. 2d). With the median dose of the original experiment, such approximations based on the median reference profile yield an average (C_av_) level in steady-state of 9.6 mg/mL of total IgG surrounded by a fluctuation from a trough (C_min_) of 7.4 mg/mL to a peak (C_max_) of 14.8 mg/mL. C_av_ levels of reference profiles at the 25^th^ and 75^th^ percentile (P25 and P75) were 5.5 and 7.3 mg/mL, respectively.

Next, we set out to transfer the measured SARS-CoV-2 neutralization potency in final containers to total IgG plasma levels – assuming no antibody-specific elimination process. We arbitrarily chose a recent, yet average, final container concentration to perform the superposition (i.e., 05/2021: 216 ± 96 IU/mL per 100 mg/mL). The resulting potential titers in plasma of PID patients over the course of 10 superpositioned dosings is shown in Fig. With a neutralization potency of ∼2 IU/mg, dosing of the original study (i.e., 4-weekly doses of 0.406 g/kg) could be approximated to yield an average steady-state level of SARS-CoV-2 neutralizing IgG (C_av_) of 20 IU/mL (C_min_: 16 IU/mL; C_max_: 32 IU/mL) (Table 1). For the same dosage but using the P25 reference profile, C_av_ in steady-state would be 15 IU/mL (C_min_: 11 IU/mL; C_max:_ 25 IU/mL) and be 22 IU/mL for the P75 reference profile (C_min_: 16 IU/mL; C_max:_ 37 IU/mL).

**Table 1:** Approximated steady-state trough (C_min_), peak (C_max_), and average (C_av_) neutralizing SARS-CoV-2 IgG-levels estimated through a superposition cascade with time-staggered dosing using the median, 25th (P25), and 75th (P75) percentile baseline-adjusted and dose-normalized reference curves. Estimates are presented for doses of 0.406 g/kg-BW, i.e. the median dose level of the reference dataset, assuming a potency of 216 IU per 100 mg IVIG/SCIG

|  | Derived from<br>the median reference profile | Derived from<br>the P25 reference profile | Derived from<br>the P75 reference profile |
| --- | --- | --- | --- |
| $C_{\text{av}}$ [IU/mL] | 20 | 15 | 22 |
| $C_{\min}$ [IU/mL] | 16 | 11 | 16 |
| $C_{\max}$ [IU/mL] | 32 | 26 | 36 |

## 4. Discussion

Immunodeficient patients may not have an efficient response to vaccination and therefore rely on prophylaxis measures not only against SARS-CoV-2 infection but also other circulating pathogens. Plasma-derived Ig is the standard of care to substitute the antibody repertoire in these patients with IVIG/SCIG manufactured from thousands of donations per batch. In the light of the pandemic, these immunodeficient patients and their health care professionals are interested in if and when a potential protection from severe infection or even SARS-CoV-2 infection might be achieved.

While neutralizing plasma antibody levels remain to be validated as a correlate of protection in clinical studies, here we reported a rapid increase of SARS-CoV-2 neutralization potency in Octapharma’s IVIG/SCIG final containers by means of ELISA and microneutralization testing against SARS-CoV-2 WT (D614G). We found a very good correlation of the ELISA data to the actual virus neutralization data, which is in line with previously published observations on the performance of the Euroimmun IgG ELISA suggesting it to be a suitable surrogate to perform donation screening and even semi-quantitatively approximate SARS-CoV-2 neutralization potency in IVIG/SCIG matrices [42–44].

The first positive IVIG/SCIG final containers were detected in 12/2020, produced partly from pre-pandemic plasma and donations obtained in 08/2020 (minimum production lead time: 5 months). In contrast to a previous report, we could not detect a relevant cross-neutralizing activity in pre-pandemic batches of IVIG/SCIG [45]. The reported COVID-19 prevalence in 07/2020 was about 1% [40], while recent reports suggest a 4.8 fold higher seroprevalence during the first wave of the pandemic [46]. In fact a recent serological survey among US blood donors showed a 3.5% seroprevalence in July 2020 [47].

The observed increase in IVIG/SCIG final container neutralization titers from 02/2021 to 05/2021 (i.e., 61 to 216 IU/mL on average), fits well to the forecast by Farcet et al., who predicted 53 to164 IU/mL for the same period of time [30]. Not factored in in this prediction was the successful initial onset of the vaccination campaign in the US. The latter which had an almost exponential increase with regards to “first-dose in” in the total US population from 01/2021 until 05/2021 is potentially being responsible for the ongoing exponential increase in 06/2021 [40]. This was also reported among blood donors where with the onset of vaccinations in 12/2020 an exponential increase was seen in overall seroprevalence to over 80% in May 2021, whilst infection-induced seroprevalence plateaued at that time [47].

Given that production of Octapharma’s IVIG/SCIG takes at least 5 months from donation until filling and with the reported exponential increase in serologically relevant exposure of the US total population from the first half-year in 2021 (inferring this to be an acceptable correlate of our donor population) suggests a further increase of neutralization capacities at least until fall 2021, while the exponential increase will likely see an inflection point and turn into a saturation phase. At which level SARS-CoV-2 neutralization titers will plateau and how stable this plateau will be over time depends on many factors: e.g., (i) the increase in vaccinees over-taking the CP donors in numbers might potentially lead to a different “on-average”, potentially more consistent contribution per plasma donation [48–50]. Our assessment of a small subset of convalescent donors (n = 313) showed that there was a marked spread in reactivity across the donations. Almost half of convalescent donations was non- or low reactive. This was more pronounced than in earlier reports, with almost a quarter of all donors as low/no responders [51]. In our analysis, only about 10% of the investigated convalescent single donations could meet actual titer demands under revised emergency use authorization (EUA) issued by FDA for CP treatment in acute COVID-19 infection [5]. However, these rare high-antibody titer donations eventually drive the titer in plasma pools. (ii) Furthermore, first case reports have been described where convalescent patients received vaccinations with mRNA vaccines boosting their neutralization titers at least 20x compared to their previous levels [52, 53]. These booster effects will likely impact potencies in individual plasma donations and subsequently in the IVIG/SCIG final containers. (iii) Even more interesting will be to see on which level anti-SARS-CoV-2 antibodies prevail in the long run, given waning of neutralization titers [54] and loss of sterile immunity on the mucosal surface of donors potentially allowing mild natural SARS-CoV-2 re-infection [55] and subsequent “natural” titer boosts. A model for this could be the detectable levels of seasonal coronaviruses in IVIG/SCIG formulations [30, 56]. Additionally, depending on public health policies, repeated boost vaccinations may also drive long-term antibody levels against SARS-CoV-2 [57].

(iv) Leaving aside the insecurity regarding the quality of individual donations, we must emphasize that the manufacture of IVIG/SCIG also involves the pooling of intermediates and donations over a collection time span of up to one year, which has the potential to average out epidemiologic trends, such as vaccination levels.

While final container neutralization potencies against SARS-CoV-2 (D614G) are strongly increasing, it is of interest how these could translate into anti-SARS-CoV-2 IgG steady-state trough levels in a patient, which is one possible, yet not validated, correlate of immune protection. Approximations of the steady-state trough levels based on a 4-weekly IVIG dosing by superposition of the median baseline-adjusted reference curve - derived from actual trial data [34] - indicate that doses of 0.4 g/kg yield a trough level of about 7.4 mg/mL IgG. These superpositioned values correspond well to previously published IVIG steady state plasma levels [58].

As a basis for our calculation, the median anti-SARS-CoV-2 neutralizing potency of 10% IVIG in 05/2021 was chosen with 216 IU/mL or 216 IU/100 mg = 2.16 IU/mg; rounded 2 IU/mg. The approximated trough levels would correspondingly be about 16 IU/mL (P25: 11 IU/mL, and P75: 16 IU/mL) – provided there are no underlying idiotype-specific elimination processes. The variation found in-between PK profiles is in our opinion less important to an actual antibody plasma trough level (Table 1), as the actual SARS-CoV-2 neutralization titer in the final container for which we have observed a substantial variation as can be seen in Fig. 2b. Likewise, the spans between the investigated time periods has more impact on the theoretical plasma trough levels than variation within PK reference profiles in [34]: a final container with the maximum neutralization potency measured (06/2021: 864 IU/mL) would result in 64 IU/mL plasma trough levels, while the average final container potency in 02/2021 (61 IU/mL) would lead to 5 IU/mL plasma trough level. To put these potential plasmatic NC50 values into context, Khoury and Cromer have recently published meta-analyses providing a first estimate of a protective level against detectable SARS-CoV-2 infection. They estimated a 50% protection level from symptomatic SARS-CoV-2 infection to be around 20% of the CP titer (54 IU/mL; 95%CI: 30-96 IU/mL) [50]. Since effects of cellular immunity induced by vaccination might confound this suggested protective measure - and patients receiving IVIG/SCIG treatment are immunologically naïve or lack a potent cellular immunity - especially for full protection, the protective threshold could be much higher. Also, it has to be kept in mind that neutralizing plasma antibodies have so far not been validated as a correlate of protection. In this light, the calculated anti-SARS-CoV-2 antibody trough plasma levels (based on the average final container potency from 05/2021: 16 IU/mL) are interpreted as sub-protective levels against infection (96 IU/mL; upper 95% CI limit).

Although the epidemiological comparison to the US total population [40] is a sub-optimal - yet easily available - indicator of vaccination and COVID-19 convalescence for the actual donor population [47] and given what we observe in individual IVIG lots, an increase in anti-SARS-CoV-2 antibody titer to meet trough levels suggested by Khoury and Cromer et al. seems possible. The current maximum NC50 value measured in IVIG/SCIG was 864 IU/mL in 06/2021 and most seroconversions have occurred only recently (keeping in mind the >5-month lead time of IVIG/SCIG production). As discussed earlier, vaccination of convalescent donors also has been described to boost immunity, rendering individual plasma contributions potentially higher in titer than early in the pandemic [52, 53]. However, production of IVIG/SCIG involves intermediate production and pooling, which offers the potential to average plasma donation periods and hence not strictly follow donors’ epidemiology. While it is prudent to speculate on the ongoing increase of titers in IVIG/SCIG final containers, the long-term perspective of anti-SARS-CoV-2 antibody levels in IVIG/SCIG remains elusive and is surely impacted by, e.g., long-term (sterile) immunity of the individual, public health policies (booster immunizations), and circulating SARS-CoV-2 variants. Furthermore, the required anti-SARS-CoV-2 titer in IVIG/SCIG has to be carefully interpreted as many uncertainties could play a role: e.g., (i) antibody levels without the back-up of cellular immunity have so far not been shown as a correlate of protection in immunocompromised patients. There are initial data that antibodies can protect against infection following recombinant monoclonal therapy in this group of patients [17]. (ii) The actual PK of anti-SARS-CoV-2 antibodies may differ from on-average IVIG PK profiles, resulting in different plasma trough levels. Here, clinical studies in IVIG/SCIG-receiving patient cohorts are needed to provide information on plasma IgG levels in these patients and correlate these with the potencies found in the infused formulations. A further limitation is that the data presented here were generated using the D614G variant close to Wuhan wildtype strain, while the pandemic is currently driven by SARS-CoV-2 variants of concern and -interest. Nevertheless, polyclonality of IVIG/SCIG could render a complete neutralization evasion of variants less likely as compared to monoclonal therapies. However, for polyclonal CP (from one donor) immune escape of variants of concern from neutralization has been shown with approximately 15x lower neutralization titers, underscoring the general notion that optimal neutralization/protection is achieved against the isolate that caused the immune response [59]. In a passive immunization setting with IVIG/SCIG, this means not only trailing the epidemiology of donors in terms of neutralizing titer but also in terms of quality, meaning which variants will be neutralized best.

## 5. Conclusion

Here, we report the onset of SARS-CoV-2 neutralizing antibodies in IVIG/SCIG final containers from 12/2020 with an exponential increase in neutralizing titers. The average values ± SD found were 02/2021: 61 ± 37 IU/mL; 04/2021: 107 ± 32 IU/mL; 05/2021: 216 ± 96 IU/mL and 06/2021: 506 ± 242 IU/mL. The maximum NC50 measured so far was 864 IU/mL.

Furthermore, we arbitrarily chose a recent, yet average, final container concentration to convert neutralization potencies of IVIG/SCIG final containers to possible trough-plasma levels in steady-state. Trough-levels of 16 IU/mL were projected based on the average neutralizing titer in 05/2021. While this value seems low, the highest detected final container potency of 864 IU/mL (maximum in latest 06/2021 time-point) would result in a theoretical trough level of 64 IU/mL.

However, which plasma levels are needed for protection against SARS-CoV-2 infection of immune-compromised patients is currently under discussion. A first estimate postulates 30-96 IU/mL with several caveats and limitations. It is therefore a matter of further clinical investigations to verify and pinpoint a protective antibody level in immunocompromised patients – without (or with reduced) the background of cellular immunity.

Despite a considerable lot-to-lot variability, a further increase based on the epidemiologic developments in the US plasma donor population seems plausible at least until fall 2021.

## Declarations

### Trademarks

Octagam^®^, Cutaquig^®^ and Panzyga^®^ are trademarks of the Octapharma group.

## Availability of data and material

Material that is still in existence and owned by Octapharma can be made available in principle for research purposes by non-profit organizations upon justified request.

WHO International Standard was obtained from NIBSC, Potters Bar, UK (see materials and methods section) [36]. All data except epidemiologic data on seroconversion generated or analyzed during this study are included in this published article and its supplementary information files.

Epidemiologic data on United States status of vaccination and COVID-19 cumulated cases were obtained from the CDC Data Tracker website [40].

## Code availability

Not applicable

## Ethics approval

Not applicable

## Consent to participate

All plasma donations analyzed in this study were obtained after informed consent of the donors. Plasma products analyzed within this study were produced from material donated after informed consent for the purpose of producing pharmaceutical products.

## Consent to publish

Not applicable

## Authors’ contributions

All authors contributed to the study conception and design. Material preparation, data collection and analysis were performed by Andreas Volk, Caroline Covini-Souris, Christian de Mey and Denis Kuehnel. The first draft of the manuscript was written by Andreas Volk and all authors commented on subsequent versions of the manuscript. All authors read and approved the final manuscript.

## Acknowledgements

The authors cordially thank Prof. Dr. Christian Drosten, Charité Berlin, Germany, for the provision of virus isolate Human 2019-nCoV ex China_BavPat1/2020_Germany ex China.

Special thanks go to Gabriele Hoch and Lisa Oberland for virus and cell culture as well as the staff of Virus and Prion Validation at Octapharma for provision of expertise and support.

The authors cordially thank Sonja Hoeller and Bernhard Rohrbacher (Global Medical and Scientific Affairs, Octapharma Pharmazeutika Produktionsgesellschaft m.b.H, Austria) for their input and critically reviewing the manuscript.

We thank Monica Byrd (Octapharma Plasma Inc., USA), Sabine Simlinger and Michael Szkutta (Octapharma Pharmazeutika Produktionsgesellschaft m.b.H Austria) for the provision of plasma donations as well as anonymized, information on non-personal records and data.

We thank the CoVIg Alliance for collaboration and exchange.

## Supplementary material

**Supp.Fig. 1.**
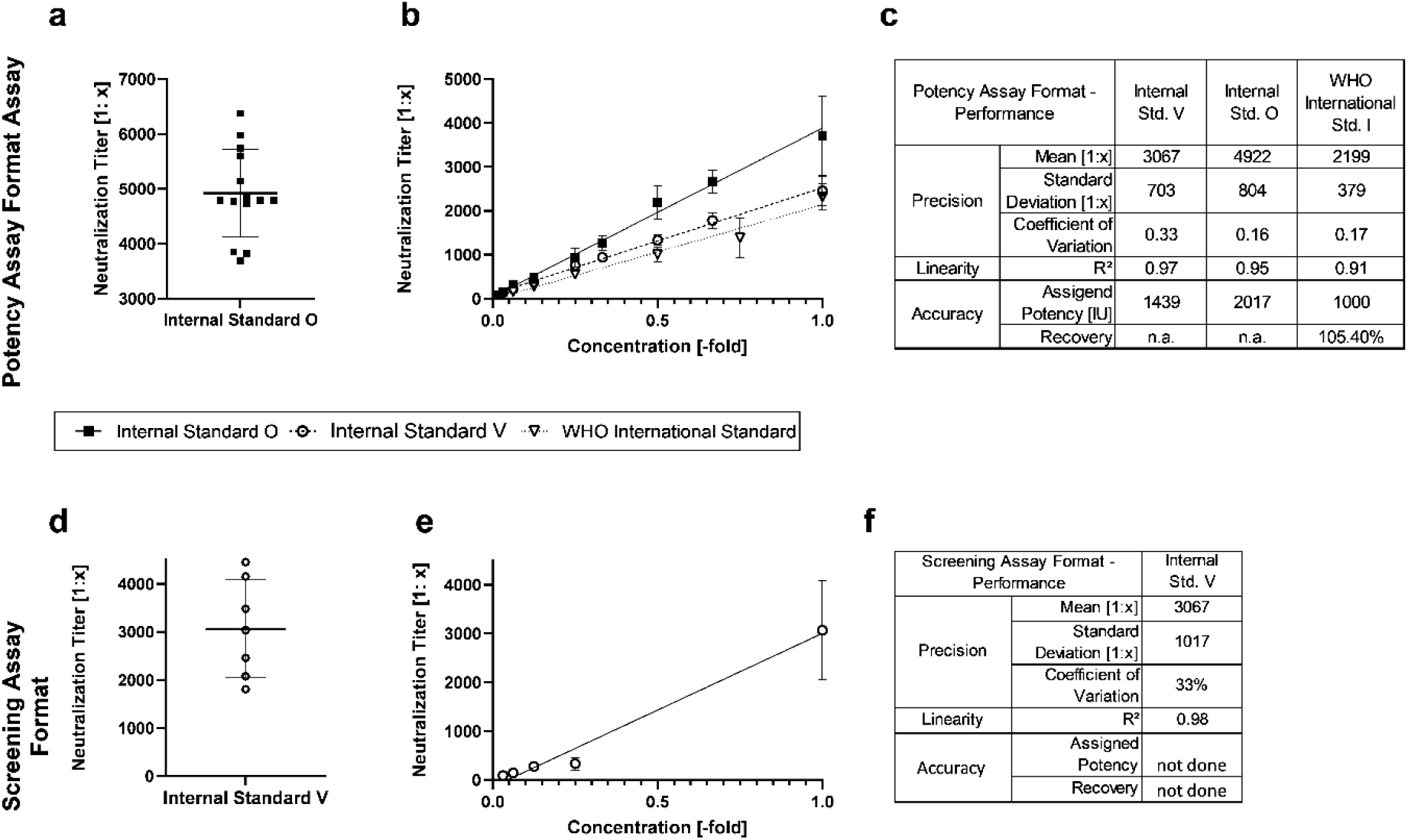
Retrospective SARS-CoV-2 microneutralization assay characteristics for potency assay format (**a-c**) and screening assay format (**d-f**). **a** and **d** intermediate precision of internal standards O and V measured as fold dilution as calibrator in each experimental run. **b** and **e** linearity data of the two assay formats. Depicted is the mean result ± SD per dilution level for each standard tested. **c** and **f** tabular results of retrospective analyses on potency and screening assay formats for precision, linearity and accuracy, where applicable

**Supp. Fig. 2:**
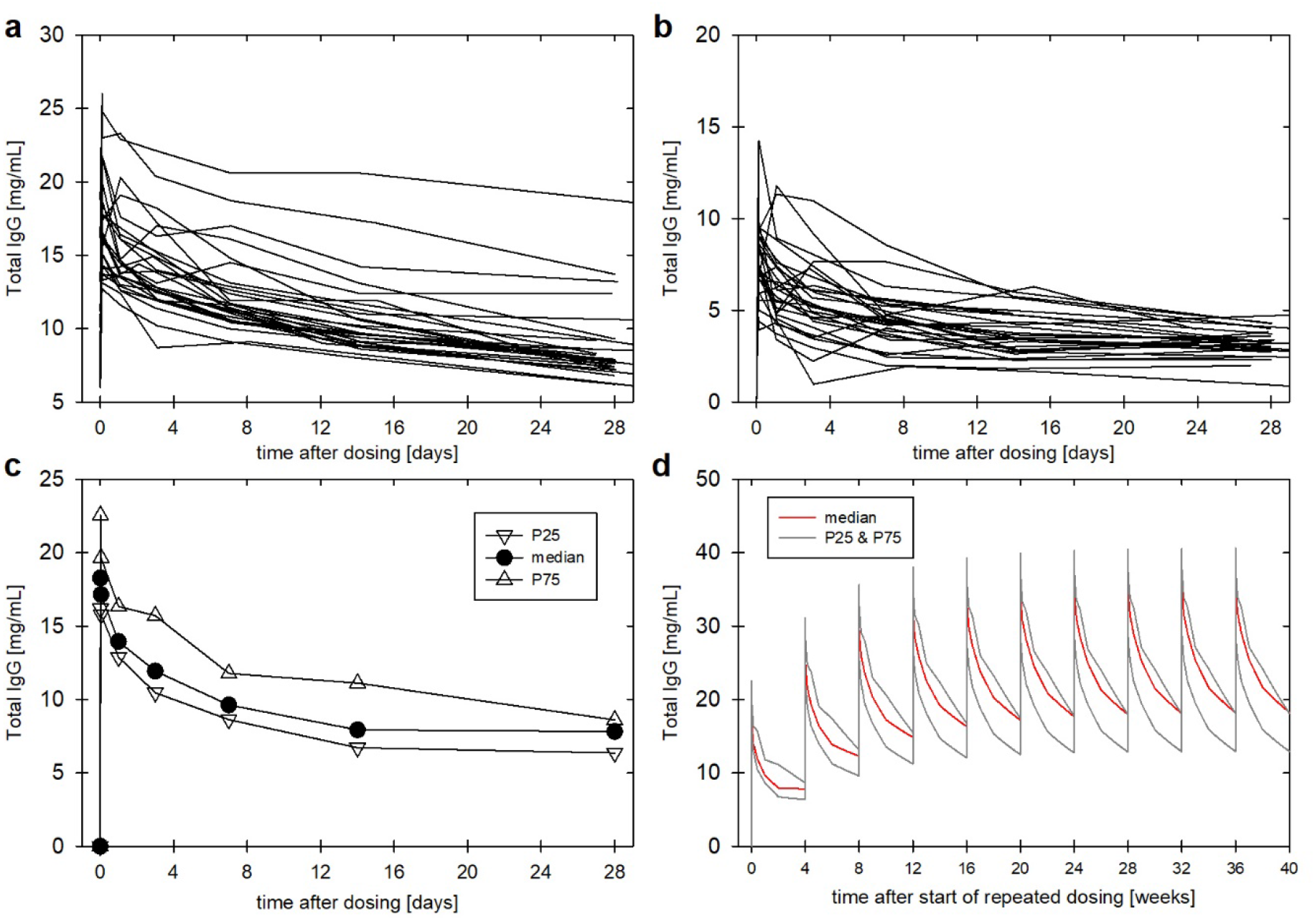
PK data processing for superposition analyses. **a** reference dataset of the observed individual time courses of untransformed IgG-levels on 4-weekly repeated dosing of IVIG (median dose: 0.406 g/kg-BW) [34]. **b** individual time courses of the baseline-adjusted IgG-levels on 4-weekly dosing of IgG. **c** reference single-dose profiles, i.e. time courses of the median, 25th (P25), and 75th (p75) percentiles of the data from panel b dose-normalized by g/kg-BW. **d** approximated time courses of the dose-normalized IgG-levels (mg/mL per g/kg-BW dosed) throughout ten 4-weekly repeated doses of IVIG derived from the median, P25 and P75 reference profiles (see panel c) using a superposition cascade with time-staggered dosing

